# Performance Limitations in Sensorimotor Control: Tradeoffs between Neural Computation and Accuracy in Tracking Fast Movements

**DOI:** 10.1101/464230

**Authors:** Shreya Saxena, Sridevi V. Sarma, Munther Dahleh

**Affiliations:** Department of Electrical Engineering and Computer Sciences, Massachusetts Institute of Technology, Cambridge MA 02139; Department of Biomedical Engineering, Johns Hopkins University, Baltimore MD 21210

## Abstract

The ability to move fast and accurately track moving objects is fundamentally constrained by the biophysics of neurons and dynamics of the muscles involved. Yet, the corresponding tradeoffs between these factors and tracking motor commands have not been rigorously quantified. We use feedback control principles to quantify performance limitations of the sensorimotor control system (SCS) to track fast periodic movements. We show that (i) linear models of the SCS fail to predict known undesirable phenomena, including skipped cycles, overshoot and undershoot, produced when tracking signals in the “fast regime”, while non-linear pulsatile control models can predict such undesirable phenomena, and (ii) tools from nonlinear control theory allow us to characterize fundamental limitations in this fast regime. Using a validated and tractable nonlinear model of the SCS, we derive an analytical upper bound on frequencies that the SCS model can reliably track before producing such undesirable phenomena as a function of the neurons’ biophysical constraints and muscle dynamics. The performance limitations derived here have important implications in sensorimotor control. For example, if primary motor cortex is compromised due to disease or damage, the theory suggests ways to manipulate muscle dynamics by adding the necessary compensatory forces using an assistive neuroprosthetic device to restore motor performance, and more importantly fast and agile movements. Just *how* one should compensate can be informed by our SCS model and the theory developed here.

## 1 Introduction

Tracking fast unpredictable movements is a valuable skill, applicable in many situations. In the animal kingdom, the context includes the action of a predator chasing its prey that is running and dodging at high speeds, for example, a cheetah chasing a gazelle. The sensorimotor control system (SCS) is responsible for such actions and its performance clearly depends on the relaying capabilities of neurons and the dynamics of muscles involved. Despite these obvious factors that set the limits on how fast an animal can track a moving object, tracking performance of the SCS and its dependence on neural relaying and muscle dynamics has not been explicitly quantified. This study aims to rigorously quantify the performance tradeoffs between accurately tracking fast moving objects, in particular periodic inputs, and the relaying capabilities of neurons subject to their biophysical constraints. The tradeoffs are also described as a function of muscle dynamics.

Tracking periodic signals is in itself an experimental paradigm that has been explored in many visuo-motor and memory tasks ^1, 2^. It has been shown in experimental settings that tracking faster and faster periodic inputs leads to a regime where subjects skip cycles because they are unable to keep up. In ^3^, monkeys performed an oculomotor task in which they were required to track periodic inputs with their eye muscles. Tracking was shown to be accurate at low frequencies, but as the frequency of the input increased, the monkeys performance deteriorated and they started to skip cycles (Fig. 1A). In ^4^, human subjects were asked to make quick periodic downward-upward motions upon hearing a periodic auditory stimulus. When the frequency of the input increased, accuracy decreased. More importantly, several undesirable phenomena including overshoot, undershoot, and skipped cycles were observed in the high frequency regime (Fig. 1B).

**Figure 1:**
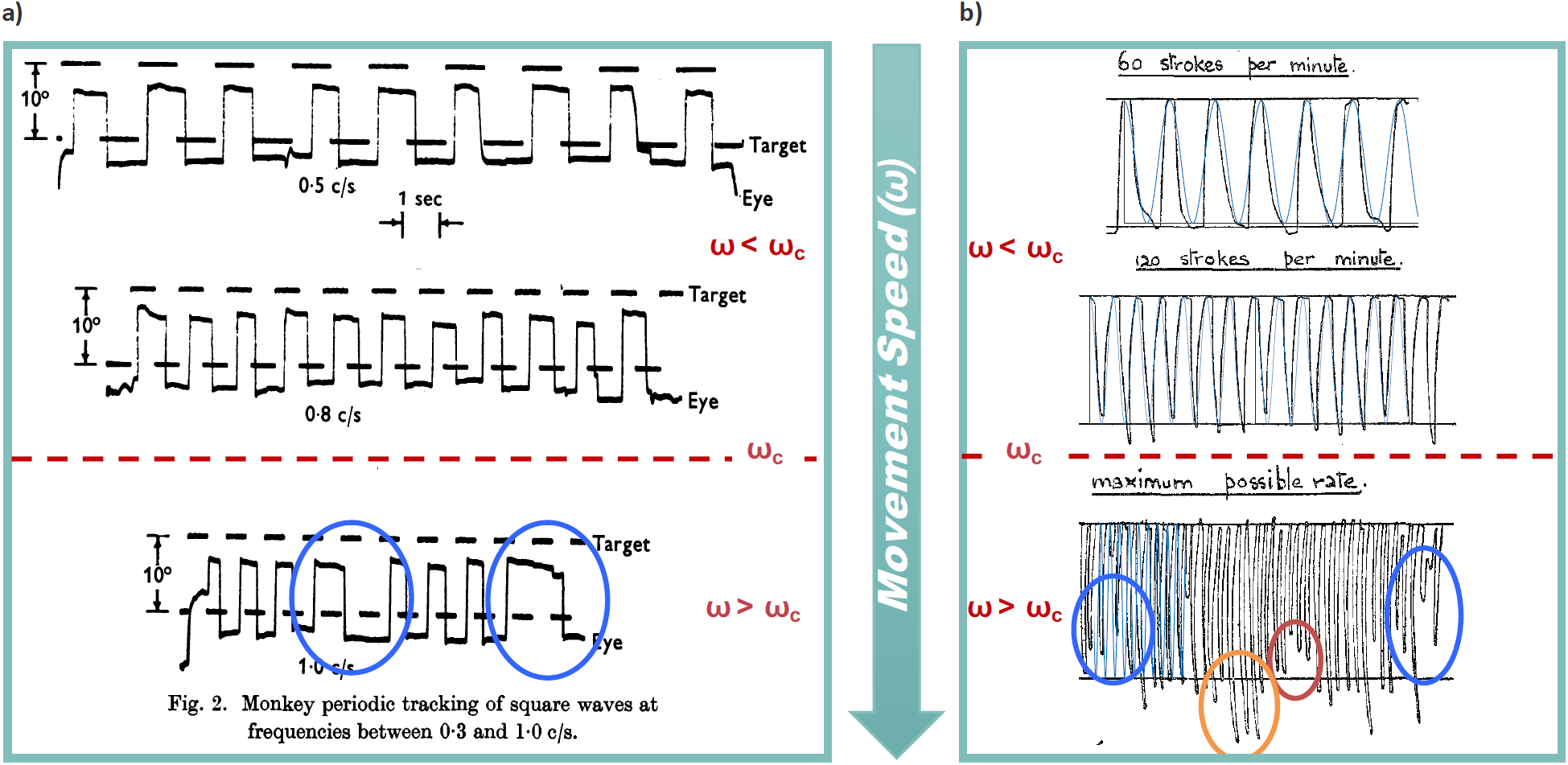
Undesirable phenomena as seen in experiments performed in a) ^3^ and b) ^4^. In both cases, fairly accurate tracking is seen in relatively low frequency reference signals (i.e. *ω* < *ω*_*c*_), whereas undesirable phenomena such as skipped cycles (blue circles), overshoot (orange circle) and undershoot (red circles) are seen in high frequency signals for *ω* > *ω*_*c*_ for some *ω*_*c*_.

We hypothesize the existence of a fundamental frequency, *ω*_*c*_, above which we observe these undesirable phenomena when tracking sinusoidal inputs. To test our hypothesis, we set out to construct a mathematical model of the SCS consisting of a feedback interconnection between the cerebrocerebellar system, alpha motor neurons and the musculoskeletal system, and then (ii) develop formal methods to prove the existence of and to compute *ω*_*c*_ as a function of our model parameters.

Although tracking in motor control has been well studied, with competing mathematical models to describe the phenomena observed ^1^, the applicability of these models in the regime of tracking *high frequency* periodic signals has been overlooked. One possible reason is due to analytical tractability of experimentally validated models. Linear models of the SCS are simple to analyze ^5^, but they fail to capture the undesirable tracking phenomena observed experimentally. For example, if you input a high frequency periodic input into a linear model, the same frequency appears at the output, but is likely attenuated. There are no skipped cycles or overshoot phenomena as observed in the SCS.

Discontinuities appearing in slow movements are a well known feature that have been studied to elucidate the junction between neuronal activity and the musculoskeletal system ^6^. Specifically, during slow movements, discontinuities in the finger position have been observed, even when subjects try hard to follow a smooth trace ^1, 7^. These discontinuities are not captured with linear models of the SCS ^1^, also discussed in Section 3. Due to this limitation of linear models, nonlinear threshold-based pulsatile control of movements was first suggested in 1947 ^8^, and since then has been in active exploration in the feedback setting ^1, 7, 9–14^.

Here, we construct an biophysically-based, analytically tractable nonlinear model of the SCS that reproduces several characteristics of neuronal and motor outputs in response to periodic inputs in both the slow and fast regimes observed in humans and primates both in health and in disease. A high level view of the model is shown in Fig. 2, while a more detailed model considered in this paper is shown in Figure 3. The critical part of our SCS model that must remain biophysically-based and nonlinear is how *alpha motor neurons* encode signals from the cerebrocerebellar system in the form of spike events.

**Figure 2:**
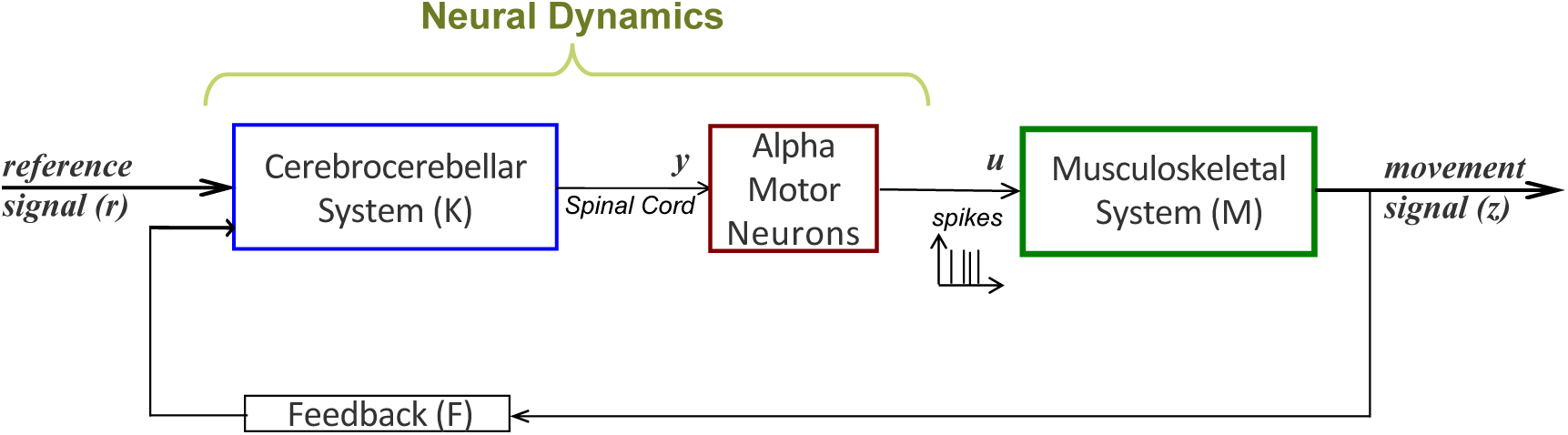
Closed loop model of the Sensorimotor Control System (SCS). The input, *r*(*t*), goes through a linear model of the cerebrocerebellar system *K*, that in turn generates a control signal, *u*(*t*), from a group of alpha motor-neurons. These motor neurons then actuate a linear model of the musculoskeletal system *M* that generates the motor output *z*(*t*), which is fed back to the cerebrocerebellar system.

**Figure 3:**
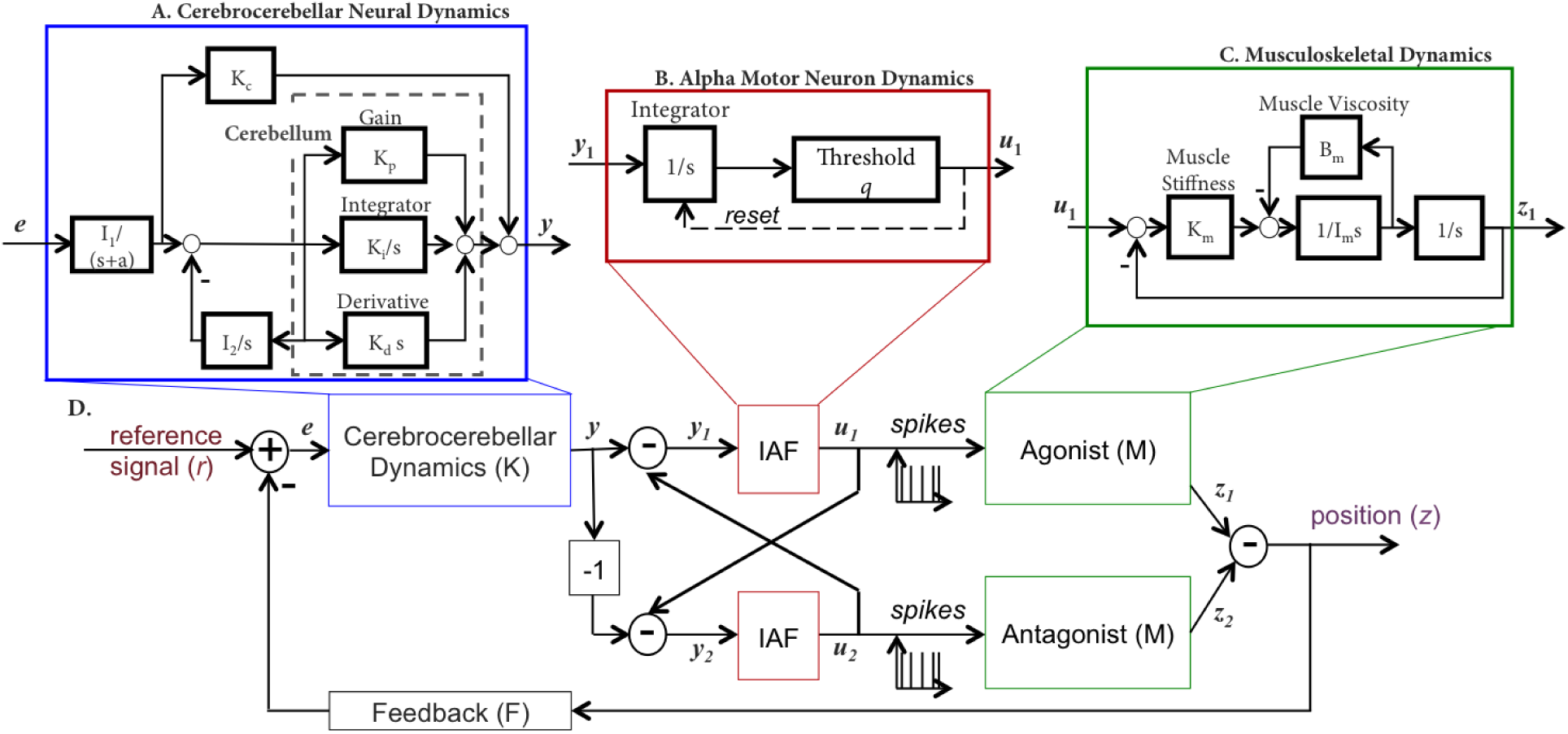
Proposed closed-loop model for movement generation. A. Details of cerebrocerebellar dynamics, adapted from ^16–18^. B. Details of the IAF Model ^19^. C. Details of the musculoskeletal system, adapted from ^20^. D. A closed-loop model of the agonist / antagonist muscles acting around a single joint.

Using our SCS model, we derive conditions for which tracking a sinusoidal input produces undesirable phenomena as a function of the parameters of the closed-loop system. Specifically, we show that for a fixed musculoskeletal system *M*, cerebrocerebellar dynamics *K*, and a group of motor neurons with spiking thresholds *q* (for activation of agonist/antagonist pair of muscles), the oscillatory input (periodic back-and-forth movement) with frequency *ω* may generate undesired skipped cycles if *ω* > *ω*_*c*_(*M, K, q*).

The results derived here make a scientific contribution in advancing our understanding of how movement speeds may be limited given the processing of the driving signal through neurons, although neuronal spiking may confer additional advantages such as energy efficiency ^15^. The characterization of *ω*_*c*_ for sinusoidal movements provides a theoretical basis for further investigation for more general classes of movements. By providing a concrete dependance on the parameters of the system, our analysis can also guide the design of therapies for movement disorders caused by a compromise in *K, q* or *M*.

## 2 Model of Sensorimotor Control System

A schematic for the SCS model is shown in Fig. 2, and a more detailed model that we consider is shown in Fig. 3. It is an internconnected feedback system, as suggested by previous models for sensorimotor control ^18, 21–24^. We assume that the SCS input is the intended voluntary movement or reference signal, *r*(*t*), that exists in some part of the brain (e.g. in parietal regions). The reference input is processed by brain structures including the sensorimotor, premotor and motor cortices, and ultimately the neurons in premotor and motor areas send spike train signals to muscles via the spinal cord. Appropriate muscles are innervated to generate a movement, *z*(*t*), as the system attempts to follow or “track” *r*(*t*). The generated movement (output of SCS) is then fed back, via proprioceptive and visual feedback, represented by a feedback gain *F*, to be processed by structures including the cerebellum and sensorimotor cortex. There are other structures involved in the generation of movements such as the basal ganglia and motor thalamus not explicitly shown.

We break down our SCS model into two general components: neural dynamics and the musculoskeletal system, as in Fig. 2. The neural dynamics consists of a) cerebrocerebellar processing of the error between intended and actual movement and b) alpha motor neuron encoding of cortical motor commands. Below, we describe each model component.

### Neural Dynamics

We take a systems approach to model neural dynamics. Specifically, we use a previously validated model of cerebrocerebellar processing ^18^ that describes how motor cortical areas integrate with cerebellar processing of errors between intended and actual kinematics. This model characterizes firing rate activity of populations of cerebellar and cortical neurons during movement, and produces an analog signal that is then sent to an integrate-and-fire (IAF) model describing the spiking responses of groups of alpha motor neurons.

The concept of *relaying capabilities* of neurons can be thought of as the density of the resulting spike trains that directly drive muscle activity: the higher the density of the spikes, the more information is flowing from the neural system to the musculoskeletal system.

### Cerebrocerebellar Dynamics

A basic cerebrocerebellar system diagram is given in Fig. 3A ^16–18^. We characterize the cerebro-cerebellar SCS component as a dynamical feedback control system (*K*) that compares the intended movement with actual movement. Much available behavioral and neurophysiological data on human and animal cerebellar motor control relates to stabilization of posture and accurate control of movements rather than explicit control of contact force or joint torques. Importantly, though, because of the uniformity of its circuitry, intra-cerebellar mechanisms are almost certainly common to both position and force control systems, especially for fast movements. This motivates our choice of modeling the cerebrocerebellar system, *K*, as belonging to a class of linear time invariant systems including proportional (*K*_*p*_), integral (*K*_*i*_) and derivative (*K*_*d*_) (PID) controllers as in ^18^.

More specifically,

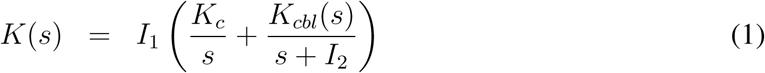

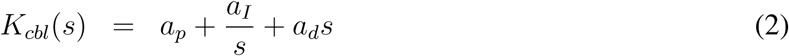

Although the cerebellar dynamics here are modeled as PID, other linear time invariant (LTI) models have been used in the past to characterize cerebellum operation ^25^; the results derived in this paper do not depend on the exact characterization for the cerebrocerebellar dynamics.

### Alpha Motor Neuron Dynamics

Alpha motor neurons are driven by the output of the cerebrocerebellar system and their activity directly drives muscles. Consequently, we maintain the spiking or pulsatile nature of alpha motor neuron control of muscles by modeling them with IAF models, which have been used to model motor neurons in both experimental and computational studies ^26, 27^.

For each joint in our muscle model described in 2, we characterize an “effective agonist group of motor units” and an “effective antagonist group of motor units”. More specifically, we assume that each group of motor units generates spikes in an IAF fashion (see Fig. 3 B). The membrane voltage of each of these groups of neurons is described as the following.

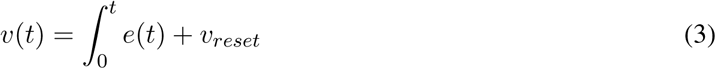

When *v*(*t*) > *q*, a weighted positive spike, *qδ*(*t*), is produced. If *v*(*t*) < −*q*, a weighted negative −*qδ*(*t*), is produced. In both cases, the membrane potential *v*(*t*) is reset to *v*_*reset*_.The collection of positive spikes 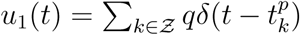 is the spike train generated collectively by neurons in the effective agonist group of motor units, and the collection of negative spikes (represented as positive action potentials in brain) 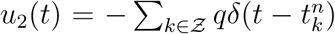 is the spike train generated collectively by neurons in the effective antagonist group of motor units. The weight of the spike is chosen to be the same as the integral, following the Henneman’s size principle ^28^. In general, if we have more than one neuron innervating the muscle, the innervation amplitude should be proportional to the activation threshold in keeping with the Henneman’s size principle. An increase in ‘relaying capability’ follows an increase in the density of the spike trains, i.e. a decrease in the spiking threshold *q*.

### Musculoskeletal Dynamics

We use a single joint model of the musculoskeletal system as in ^20^ (Fig. 3C), though our methods easily extend to multiple joint systems. The transfer function of this model from the neural activity (*u*) to the limb output (*z*) is the following.

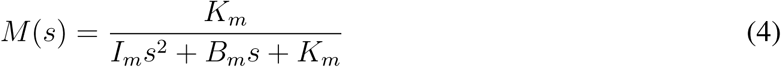

As shown in Fig. 3C, *K*_*m*_ corresponds to the net stiffness of all muscles acting around the joint, as determined by the level of agonist/antagonist coactivation, *B*_*m*_ is the net viscosity, and *I*_*m*_ is the inertia. This model is taken from the class of “equilibrium-point models” for motor control ^20^. Under this class of models, the mechanical properties of muscles and the myotactic reflexes generate equilibrium positions for the limb. If the limb is displaced from rest position, the spring-like properties of the muscles generate the appropriate restoring torques to return the limb to rest (equilibrium). These models are in reasonable agreement with experimental data, yet are simple enough to analyze. More information about the musculoskeletal model is provided in the Supplementary Information.

## 3 Results

In this section, we first demonstrate how our closed-loop SCS model qualitatively reproduces fundamental properties of alpha motor neuron activity and musculoskeletal responses during movement generation. We also show the qualitative reproduction of responses to several types of reference signals observed in both healthy subjects and cerebellar patients. We then show that a linear SCS model would not suffice to reproduce the phenomenon of skipped cycles for higher frequencies. Finally, we derive the fundamental frequency, *ω*_*c*_ as a function of model parameters, and describe its sensitivity to SCS model parameters.

### SCS Model Reproduces Observed Phenomena in Health and in Disease

he SCS model introduced in Section 2 reproduces observed phenomena by the SCS, as shown in Fig. 4. Fig. 4a illustrates how motor neurons may produce plateau potentials, resulting in self-sustained firing, providing a mechanism for translating short-lasting synaptic inputs into long-lasting motor output. This self-sustained oscillatory behavior is a key property observed in more complex models of neural behavior ^32^. Fig. 4b shows how during voluntary contraction of a muscle group, agonist and antagonist muscles can both be active with a specific activation pattern (agonist first, then antagonist, and then agonist again at a lower level). This concurrent activation of agonist and antagonist muscles is referred to commonly as a triphasic pattern of muscle contraction during rapid limb movement and is captured by our model. The model reproduces spiking patterns during slow periodic movements, as shown in Fig. 4c, in fact also capturing the discontinuities during slow movements.

**Figure 4:**
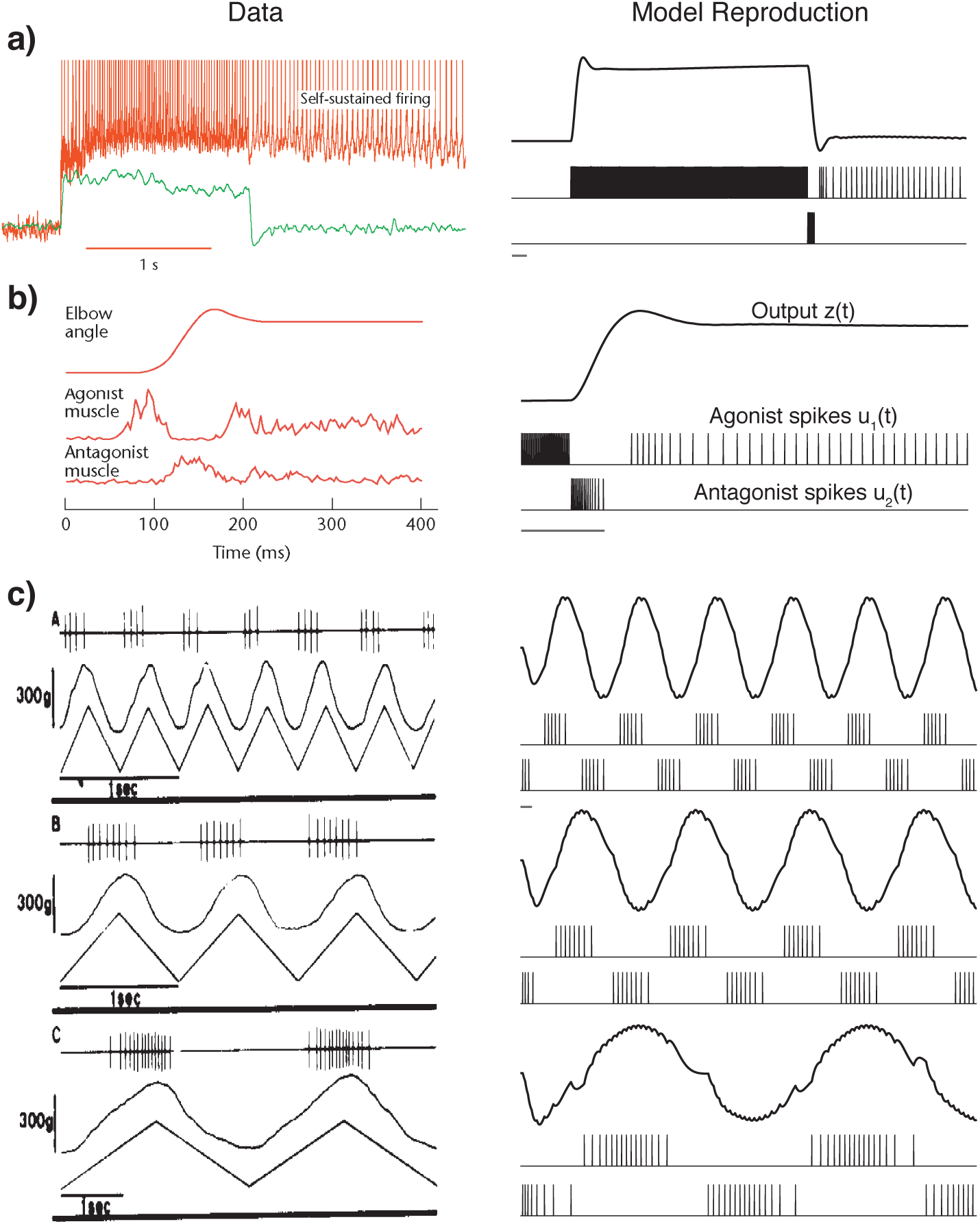
Model reproduces known experimental properties of alpha motor neuron activity. In all these subparts, experimental data from literature is to the left, and simulations using our model is to the right. Note that these are qualitative reproductions. a) self-sustained oscillations ^29^. b) triphasic pattern of muscle contraction during rapid limb movement ^30^. c) The match between spiking patterns for periodic movements in ^31^ (Left), and our simulation results (Right), all for low movement speeds, i.e. *ω* < *ω*_*c*_. The grey bar at the bottom of each plot represents 0.5 seconds.

The SCS model also reproduces movement responses to some elemental reference inputs in both health and cerebellar disease. Figures 5a and 5b compare model responses to step and pulse inputs to responses of human subjects to the same input types in health ^35^ and in cerebellar dysfunction ^33, 34^. The position signals between model and experimental data qualitatively match very well. We model cerebellar degeneration by decreasing the gain parameters of the cerebellar model (*K*_*d*_, *K*_*p*_, *K*_*i*_ and *I*_2_) by 60% as well as increasing the muscle stiffness *K*_*m*_ by 4 times, to simulate the typical compensation strategy displayed by patients ^17^. Cerebellar patients are known to display dysmetria, specifically hypermetria (or significant overshoot), as well as oscillatory activity past the end of the movement. We see this effect in the model simulations in Figs. 5a,b.

**Figure 5:**
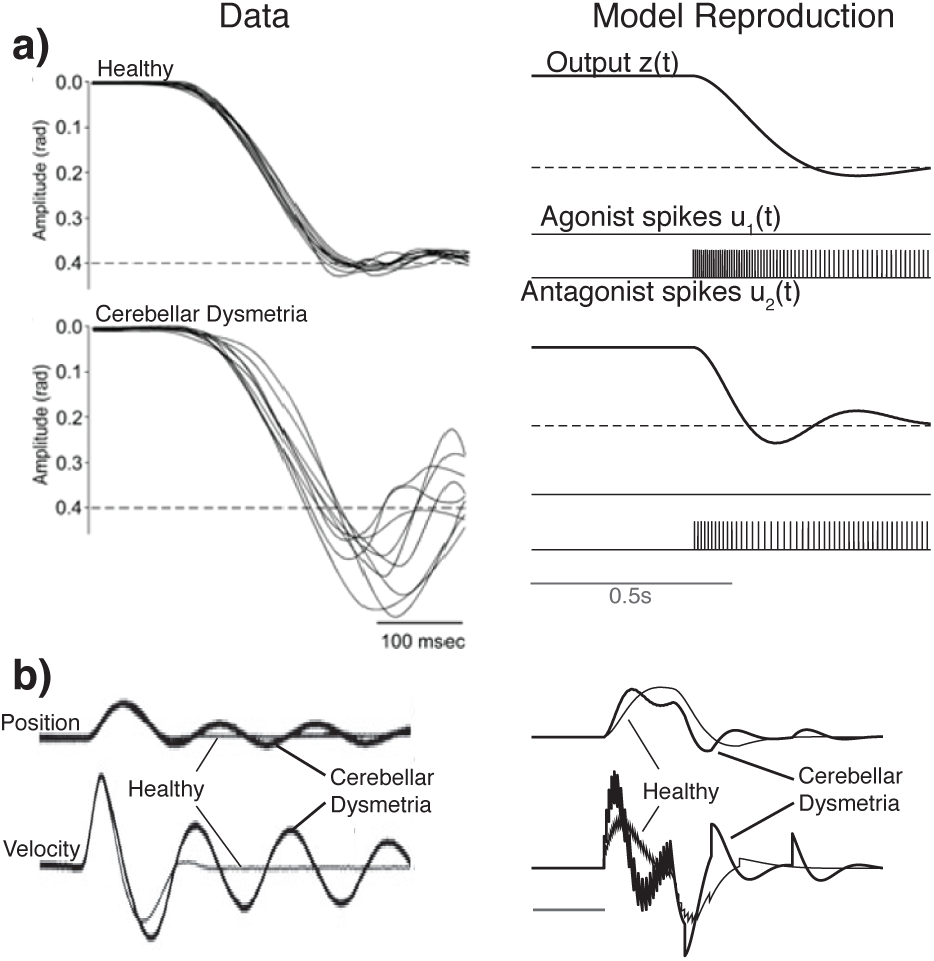
Model reproduces responses to elemental signals in health and during cerebellar disease. a) Step Movement. Data for healthy individuals and for patients with cerebellar disease ^33^, b) Short Pulse. Data for healthy individuals and for patients with cerebellar disease ^34^. Note that these are qualitative reproductions.

### Linear Time Invariant SCS Model does not display skipped cycles

The presence of skipped cycles is perhaps the most important feature observed while tracking fast sinusoidal inputs, and this phenomena (a) cannot be modeled using linear time invariant (LTI) systems, and (b) cannot be remedied in open loop using a linear system at the output of the movement. If all the components of the feedback system are linear, the steady-state output only contains the frequency present in the input signal, albeit with an amplitude and phase that depends on the gains of the individual systems. Specifically, we consider *r*(*t*) = *R* sin(2*πωt* + *ϕ*), and we denote the closed loop system from *r* to *z* as *H*. If the closed loop contains *K, M* and *F* LTI as above, and does not contain the IAF nonlinearity, 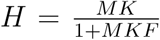 Using principles in linear dynamical systems, we can show that for any LTI *H*, the steady state response of the system can be written as the following ^36^.

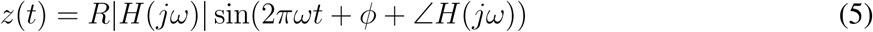

Note that *z*(*t*) is confined to oscillate at the input frequency *ω*, and thus an LTI model cannot produce either discontinuities in the movement or skipped cycles.

Moreover, the IAF nonlinearity acts in such a way as to not transmit information if the amplitude of the integral of the signal *y* falls between −*q* and *q*. This inherently ties together the amplitude of the integral of *y* and the frequency of the desired movement *r*, and leads to the conditions derived in the next section for the feedback system to reliably transmit the periodic signal. Moreover, the nonlinearity in the IAF is a thresholding and reset nonlinearity, which can be approximated by an identity function below the limiting frequency of *ω*_*c*_ ^37^.

### Spiking SCS Model Enables Analytical Derivation of *ω*_*c*_

In the SCS model as in Fig. 3, we see in Fig. 7 that our SCS model is able to maintain good tracking performance to a sinusoidal input until the input frequency is larger than the fundamental frequency *ω*_*c*_. After this input frequency, the model displays undesirable phenomena seen in experimental data, including (i) undershoot, (ii) overshoot, and (iii) skipped cycles. Both undershoot and overshoot are phenomena accompanying fast movements in general, i.e., tolerating lower accuracy while reaching for a target, and then correcting the movement when close to the target. In a repetitive movement, however, the movement is not corrected for the duration of several cycles, as seen in Fig. 1b. Here, we derive a fundamental frequency, *ω*_*c*_, for which skipped cycles are generated when tracking a sinusoidal input whose frequency is higher than *ω*_*c*_. More importantly, we compute how *ω*_*c*_ depends on our SCS model parameters. Intuitively, we know that skipped cycles are observed as the density of spikes decreases. In order to track the sinusoidal input, the alpha motor neurons must generate a series of positive spikes to track the increasing part of the sinusoidal reference input, which are then followed by a series of negative spikes to track the decreasing part of the sinusoidal reference input. To avoid skipped cycles, we need the ‘spike period’, i.e. the maximum length of time between the *sequence* of positive spikes to the *sequence* of negative spikes and back, to be less than of equal to the period of the sinusoidal input *r*. To calculate the dependence of the presence of skipped cycles on system parameters, we build a switching map from one spike to the next, and analytically determine the switching times. We can then determine whether the spike period is more than the period of the driving oscillation for some initial conditions of the system.

Briefly, let the positive and negative spike times be defined as 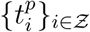 and 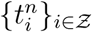 respectively. Let 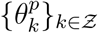 be defined as the time between the first positive spike in a sequence of positive spikes and the first subsequent negative spike, and similarly for 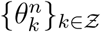. See Fig. 6.

**Figure 6:**
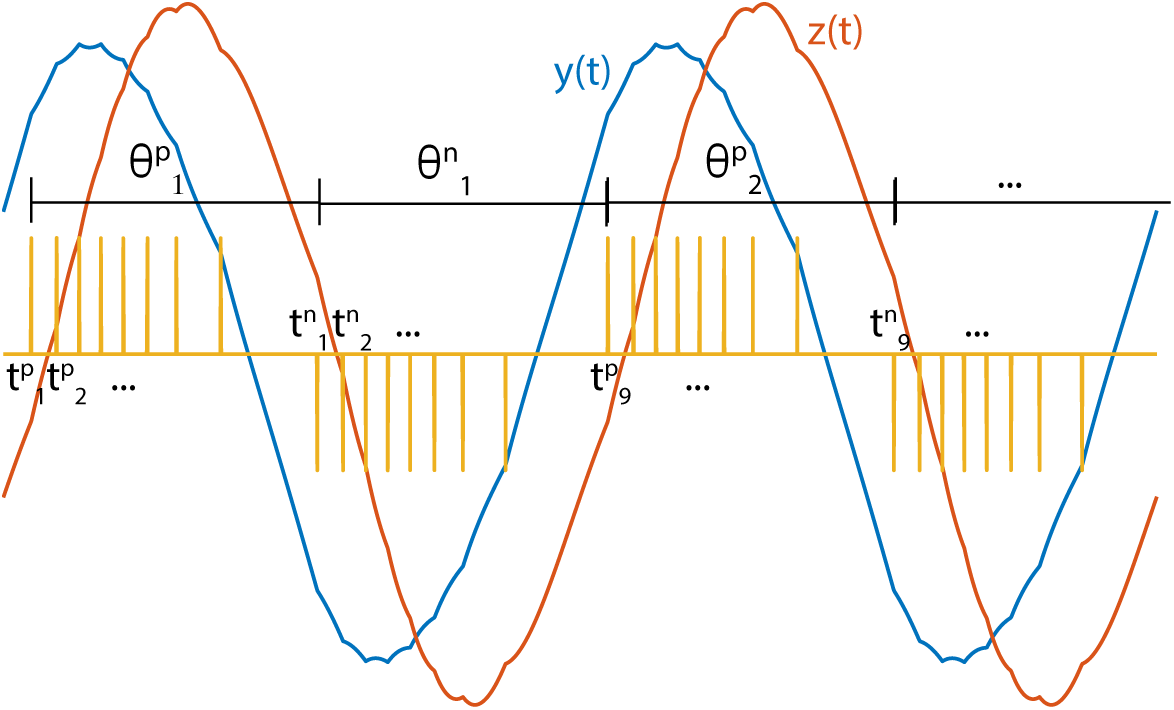
A graphical illustration of positive spike times 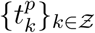, negative spike times 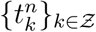 and spike period max 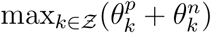

**Figure 7:**
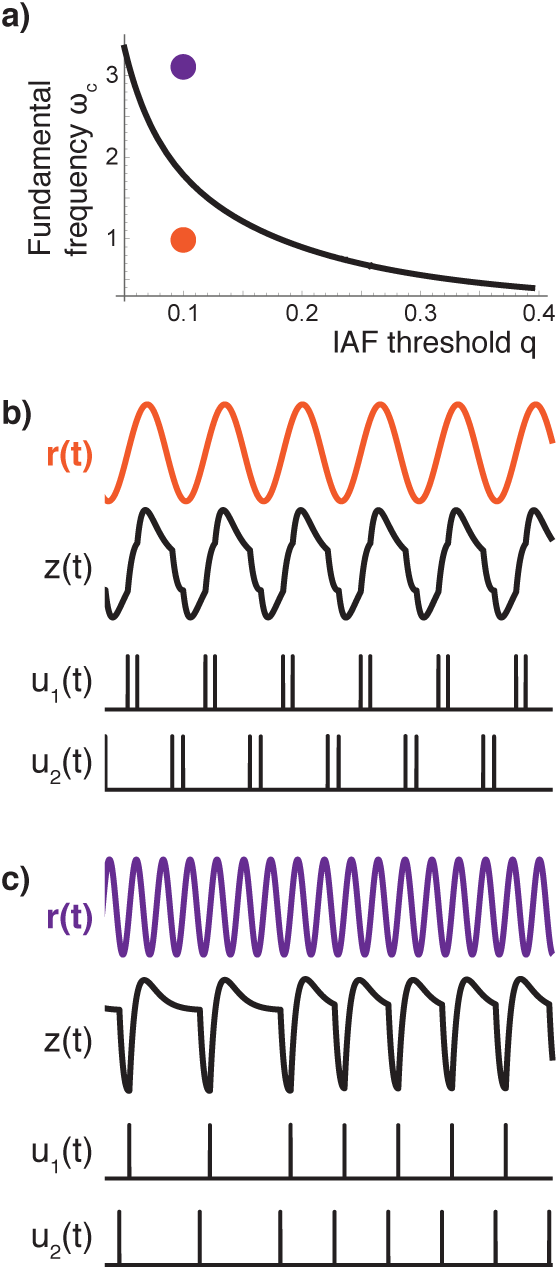
a) *ω*_*c*_ for *R* = 1, *K* = 1, *F* = 1, for varying *q*. b) *q* = 0.1, *ω* = 1. c) *q* = 0.1, *ω* = 3.18.

### Conjecture

Let *r*(*t*) = *R* sin(2*πωt* + *ϕ*), and the closed loop system as in Fig. 3D. For any given parameter values, there exists a cutoff frequency, *ω*_*c*_, such that for all *ω* > *ω*_*c*_, there exist initial conditions which lead to skipped cycles.

We first define *A*_*k*_ and *t*_*k*_ as *z* at the time of switching and the switching time, respectively. The system can be entirely described by the state *A*_*k*_ and time of switching *t*_*k*_. The switching map **Π** given a sequence of *q*^l^*s*, i.e. {±*q*, ±*q, …*} for the state and spike times is as below.

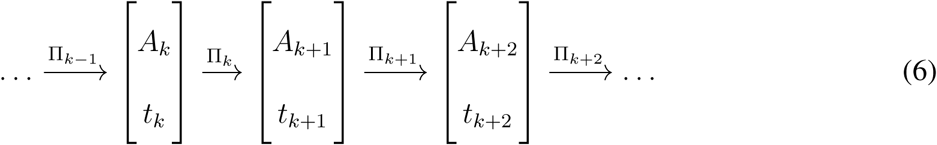

The equations for calculating Π_*k*_, i.e. calculating *A*_*k*+1_ and *t*_*k*+1_ as a function of *A*_*k*_ and *t*_*k*_, for a simple SCS model are provided in Section 6, and are easily extendable to higher order models.

In general, this system can have multiple positive spikes in a row and multiple negative spikes in a row. We define 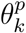 and 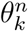 recursively as the following (see also Fig. 6).

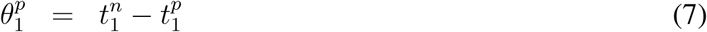

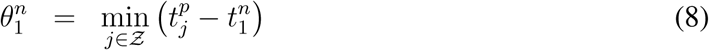

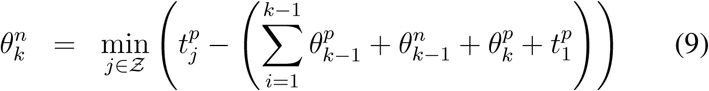

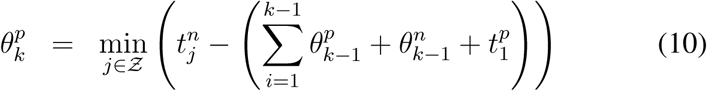

Note that for the spike train to be periodic with at least frequency *ω*, max 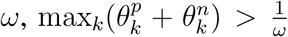. If the condition max 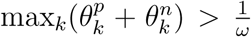 is not met, we have skipped cycles at the output *z*. Thus, we can find the dependence of 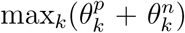 on *q* for a given system with a given sinusoidal reference signal *r*, as detailed in Section 6. In this Section, we also numerically show the conjecture for specific parameter values.

Fig. 7 shows *ω*_*c*_ as a function of *q* given a specific *M, F* and *K*. As expected, as the spiking density increases (*q* decreases), more information about the cerebrocerebellar output is transmitted to the muscles, and we see that the tracking performance increases (*ω*_*c*_ increases).

In Fig. 8, we plot the explicit dependence of the fundamental frequency on the different system parameters.

**Figure 8:**
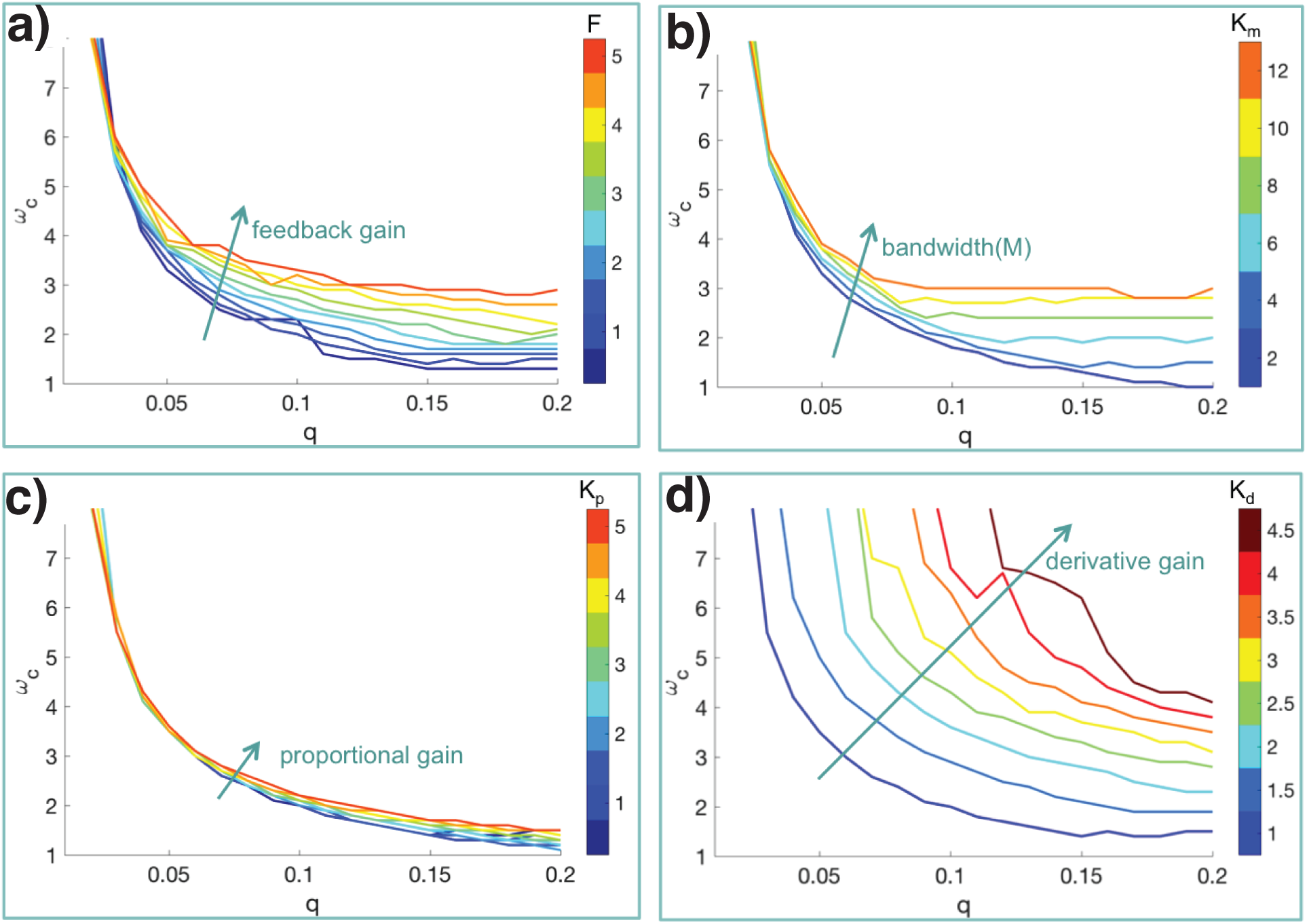
Dependence of the fundamental frequency *ω*_*c*_ on system parameters, with *R* = 1, and *K, F, M* as in Fig. 3. a) Varying the feedback gain *F*. b) Varying the muscle stiffness *K*_*M*_. c) Varying the cerebellar proportional gain *K*_*p*_. d) Varying the cerebellar derivative gain *K*_*d*_. Unless varied, parameters are as in Table 1.

We observe that as the feedback gain, *F*, or as the bandwidth of the musculoskeletal system *M* increases, we have a larger *ω*_*c*_ for all values of *q*. Moreover, as the cerebellar proportional gain (*K*_*p*_) increases, *ω*_*c*_ increases, but not significantly. However, as the cerebellar derivative gain (*K*_*d*_) increases, *ω*_*c*_ increases drastically, showing the large degree of sensitivity to *K*_*d*_. The derivative action is hypothesized to take place in the dentate neurons of the cerebellum ^17^, which are known to be important for rapidly implementing a desired action ^38, 39^. This feature is corroborated in Fig. 8d.

## 4 Discussion

Humans can make flexible and fairly fast movements, however damage to various parts of the central nervous system can limit motor performance. Whether damage is caused by stroke, multiple sclerosis, amyotrophic lateral sclerosis, or spinal cord injury, weakness (paresis) occurs routinely after injury to the primary motor cortex (M1) or its output to the spinal cord ^40^. This weakness is known to be associated with reduced voluntary recruitment of motor units in the spinal cord, both in terms of the number of motor units recruited and the firing rates they achieve ^41–45^. Given that weakness translates to deficient production of muscular forces that are used to accelerate the limbs, we expect weakness to limit fast movements, with progressively more reduction of neural activity being associated with a deceleration of voluntary movements ^40^. However, to the best of our knowledge, this is the first time that a tradeoff between relaying ability of neurons and motor tracking accuracy have been quantified. Although a “speed-accuracy” tradeoff, otherwise known as Fitts’ Law ^46^, has long been investigated for the dependence of target error on the speed of reaching to a stationary target, these tradeoffs do not address the tracking of a moving target, and the neural underpinnings have also not been conclusively determined ^6^.

Our modeling framework provides us with an explicit dependence of the emergence of undesirable phenomena (skipped cycles) as a function of *q*, the parameter that directly dictates the density of spikes (neural computing), the musculoskeletal system *M*, and cerebrocerebellar dynamics *K*. Specifically, we show that for a fixed musculoskeletal system *M*, cerebrocerebellar dynamics *K*, and a pair of motor neurons with thresholds *q* (for activation of agonist/antagonist pair of muscles), the oscillation input (periodic back-and-forth movement) with frequency *ω* may generate undesired skipped cycles if *ω* is ‘too large’ and/or the density of spikes (related to *q*) is ‘too small’. Such undesirable phenomena are consistent with symptoms observed in patients with movement disorders, and finger tracking experiments wherein subjects are trained to follow a trace back and forth quickly. Here, we address the fundamental frequency past which a sinusoidal periodic movement is no longer well tracked. Notably, this fundamental frequency may serve as a guide for movements at all frequencies, and may point towards the bandwidth of the SCS model being limited to a similar value as *ω*_*c*_. We plan to address this in future work.

The formulation in this paper also *quantifies* the increase in maximum speed gained due to the system parameters. This can help us compensate in limbs by providing either electrical stimulation directly to the muscles, or using an exoskeletal device that can provide extra assistence to the limb. An example of these ideas is presented in Fig. 9. Here, the block for neural activity contains the cerebrocerebellar model as well as the IAF block with the threshold parameter *q* as earlier. Let *ω*_*c*_ as a function of *q* for the normal regime be given by the blue curve in Fig. 9B, and let the neural activity be compromised such that *q* increases (this could correspond to a damage directly in the alpha motor neurons or the spinal cord). Decreased neural activation or sparser spikes is seen as a key symptom of general motor impairment, and we are directly modeling that effect as a decrease in the threshold to spike (recall that this is a systems-level model of neural computing, as detailed in Section 2)). In order to restore functioning, we can add a compensator as in Fig. 9A, which measures the output of the musculoskeletal system, and in turn stimulates the musculoskeletal system in feedback (here, *C* = 10 and thus 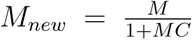). If we can further drive the output of the musculoskeletal system up by the same amount, i.e. *z*_*new*_ = 10*z*, then we can drive up the operating curve for *ω*_*c*_ to the red curve shown in Fig. 9B. This suggests that it in theory one can compensate for damage to the neural system using reinforcements, for example via neuroprosthetics at the musculoskeletal end. While this is shown in an ad-hoc manner here, the framework in this paper allows us to quantitatively design compensators, *C*, given SCS parameters, i.e. *R, q, K, F*, and *M* to meet specific control objectives. Specifically, one can design a compensator *C* with the following objective.

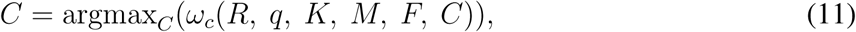

were *ω*_*c*_ can be calculated as a function of *q* and the rest of the SCS parameters as formulated above.

**Figure 9:**
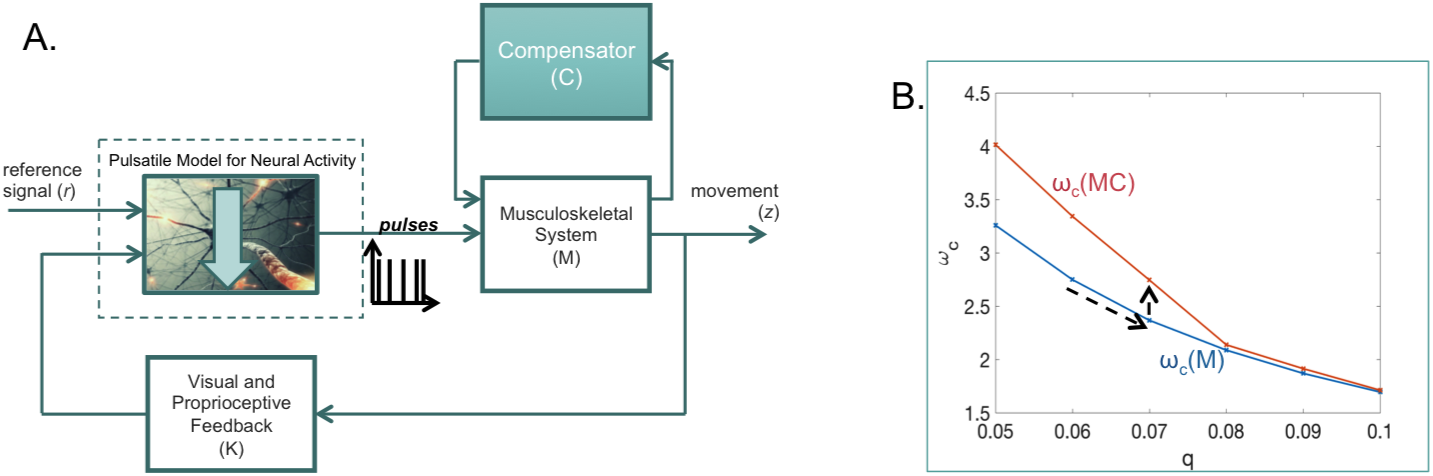
Design of a compensator to restore functioning of the SCS in the presence of a compromised neural system; more details in the text.

A critical component in our SCS model that captures undesirable tracking phenomena in the fast regime is the pulsatile control implemented by alpha motor neurons. Pulsatile constrol of movement has provided us with insights in the ‘slow’ regime, but has rarely been analyzed for limitations in tracking fast movements.

Our SCS model can reproduce experimentally known properties of motor neurons and responses to step, pulse and sinusoidal signals. However, other threshold-based pulsatile models for neural control can be readily used in the theoretical framework developed here. Furthermore, multi-joint muscle models can easily be handled by our framework as adding joints simply adds dimensionality to *M*. Our framework can also be augmented to allow incorporation of transmission delays incurred by the spinal cord. Small delays can be approximated by first order linear systems (Pade approximations as in ^36^) while larger delays in feedback interconnections can be modeled as structured uncertainty. Finally, this model allows for robustness to modeling errors while building compensators, using standard methods as detailed in ^47^.

## 6 Appendix

*A*_*k*_ **and** *t*_*k*_ **for a simple SCS model** We detail *A*_*k*_ and *t*_*k*_ here for a simple SCS model, where *K* = 1, 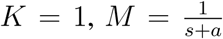, and *F* = 1, for some *a* > 0. We first describe the dynamics of *z*(*t*) for any *t* between two spikes *t*_*k*_ and *t*_*k*+1_, i.e. *t*_*k*_ ≤ *t* < *t*_*k*+1_ as the following. Specifically, 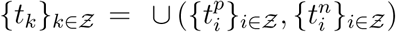

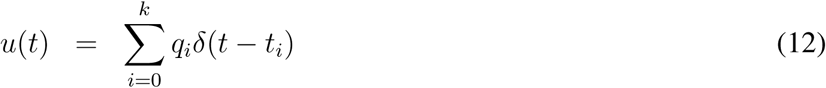

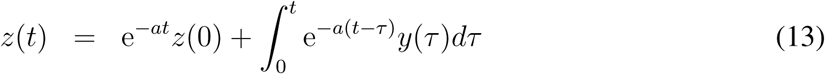

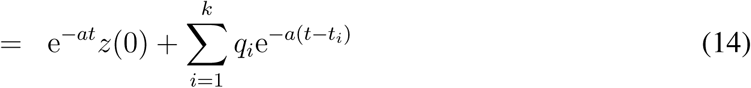

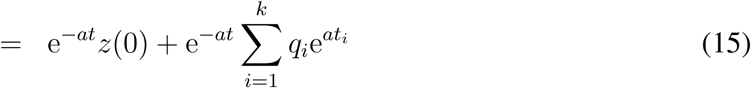

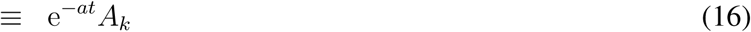

Here,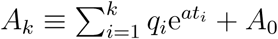, where *A*_0_ = *z*(0), and we assume that *t*_0_ = 0.

At all times, *y*(*t*) = *z*(*t*) − *r*(*t*), and the value of 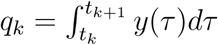.Thus, given a sequence of {*q*_*k*_}_*k*∈Z_, where *q*_*k*_ ∈ {*q*, −*q*}, we have the following condition for the dynamics.

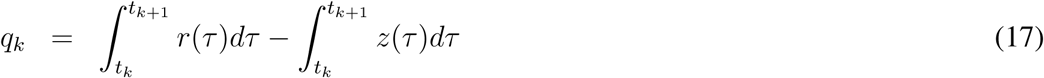

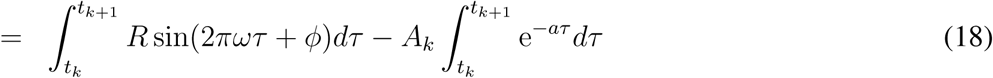

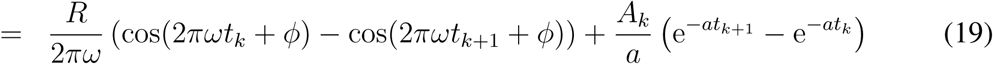

Thus, for the simple model, *A*_*k*+1_ and *t*_*k*+1_ are such that:

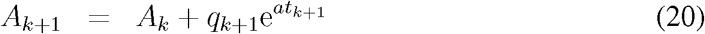

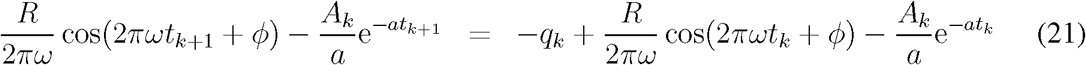

### Numerical Solution for general *K* and *M*

Due to the transcendental nature for the equations to find *A*_*k*+1_ and *t*_*k*+1_ (for example, as in Equations 20 and 21), a numerical solution is required. This is obtained by forward marching in time using a suitably chosen time step, for each set of initial conditions *z*(0) and *ϕ*. We perform the numerical solution for 20 different randomly chosen values of *z*(0) and 12 different values of *ϕ*, and take the maximum 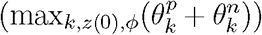 of these, hereafter written as 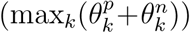 for short. In Fig. 10, we show *ω* 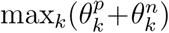 as a function of *q* for several values of *ω* for two sets of specific system parameters; one for 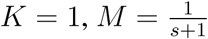, and *F* = 1, and the second for 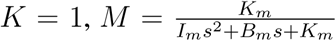, with *K*_*m*_ = 1, *I*_*m*_ = 1, *B*_*m*_ = 1, *R* = 1 and several values of *ω*. We see that for both cases, as we increase *q*, we hit the regime termed the undesirable regime, namely with skipped cycles, with 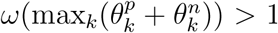. Moreover, for any fixed value for *q*, we see that there exists a frequency *ω*_*c*_, such that for *ω* > *ω*_*c*_, we can find initial conditions that lead to skipped cycles. Thus, we show the conjecture for specific systems. More details for general *K* and *M*, as well as approximation methods for determining *ω*_*c*_, are discussed in ^37^.

**Figure 10:**
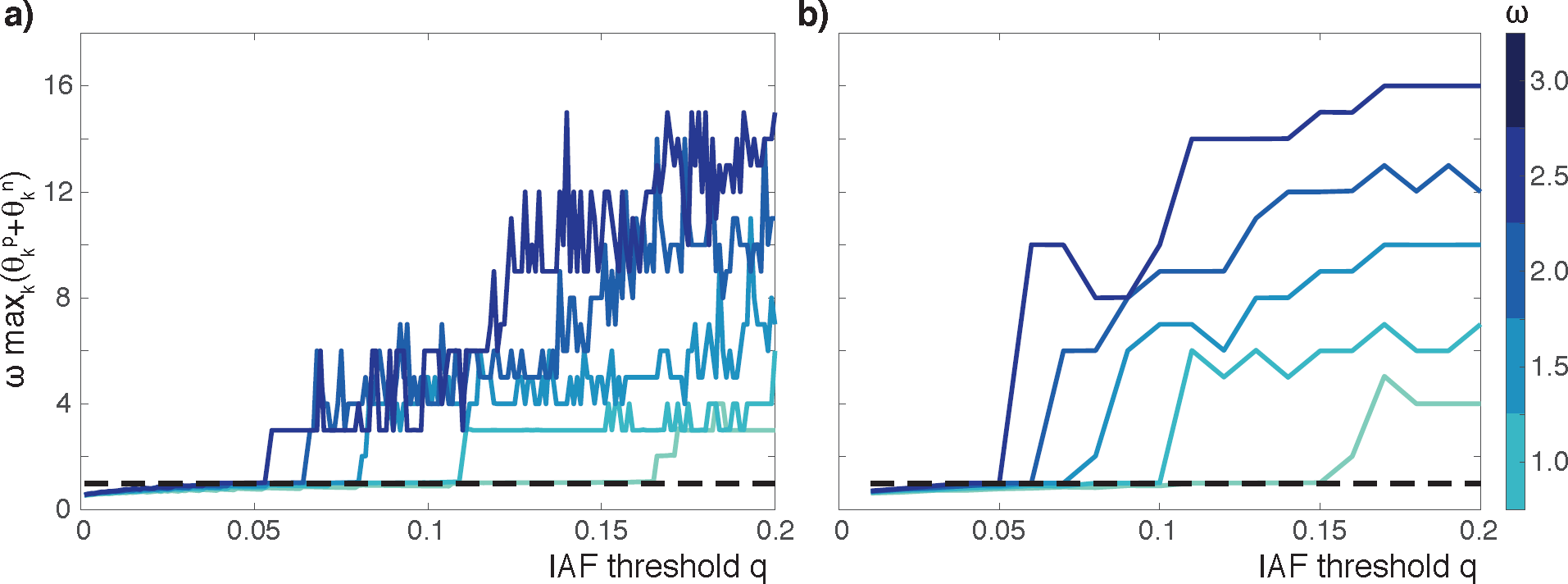
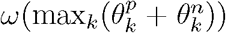, for several values of *ω* and *q*, for two different systems. If this is greater than 1, the system displays skipped cycles. 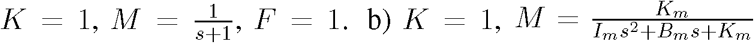, with *K*_*m*_ = 1, *I*_*m*_ = 1, *B*_*m*_ = 1. The systems were solved forward in time for multiple values of initial conditions *z*(0) and *ϕ*.

### Musculoskeletal Dynamics

Considerable work has been done on modeling the forward dynamics of the arm ^48^. In principle, models can be very complex taking into account multiple muscles ^49^, their geometries of origin and insertion ^49^, effective moment arms ^49^, activation dynamics ^48, 50, 51^, nonlinear force-velocity relations ^50, 51^, skeletal mass distributions ^48^, and spindle behavior ^52^.

As this study requires analytical tractability of the SCS model, we use a simplified model of the musculoskeletal system of a single joint as in ^20^. In this model, the muscle acts like a spring for passive displacements. This formulation is consistent with alpha-gamma coactivation that maintains the sensitivity of the muscle model to multiple inputs; deafferented monkey experiments in which the proprioceptive feedback to the spinal cord is compromised; as well as reflex function of the limbs. Analysis can also be carried out using Hill-type models for a single muscle ^53^; however, various combinations of these Hill-type muscle models would be needed to model a single joint.

---

**Table 1:** Table showing the values used for model parameters. The corresponding model
is shown in Fig. 3.

| Parameter | $K_c$ | $K_p$ | $K_i$ | $K_d$ | $I_1$ | $I_2$ | $K_a$ | $K_m$ | $I_m$ | $B_m$ | $K_f$ |
| --- | --- | --- | --- | --- | --- | --- | --- | --- | --- | --- | --- |
| Value | 0.5 | 0.5 | 0.1 | 1 | 1 | 0.2 | 1 | 4 | 0.1 | 1 | 1 |

